# Principal Component Analysis and its Generalizations for any type Sequence (PCA-SEQ)

**DOI:** 10.1101/535112

**Authors:** V.M. Efimov, K.V. Efimov, V.Y. Kovaleva

**Affiliations:** Institute of Cytology and Genetics SB RAS, Novosibirsk, Russia; Institute of Systematics and Ecology of Animals SB RAS, Novosibirsk, Russia; Novosibirsk State University, Novosibirsk, Russia; Tomsk State University, Tomsk, Russia; Moscow Institute of Physics and Technology (State University), Moscow, Russia

**Keywords:** time series, SVD, PCA, PCo, MDS, SSA, molecular sequences, p-distance

## Abstract

In the 40s of the last century, Karhunen and Loève proposed a method for processing of one-dimensional numeric time series by converting it into a multidimensional by shifts. In fact, a one-dimensional number series was decomposed into several orthogonal time series. This method has many times been independently developed and applied in practice under various names (EOF, SSA, Caterpillar, etc.). Nowadays, the name SSA (the Singular Spectral Analysis) is most often used. It turned out that it is universal, applicable to any time series without requiring stationary assumptions, automatically decomposes time series into a trend, cyclic components and noise. By the beginning of the 1980s Takens showed that for a dynamical system such a method makes it possible to obtain an attractor from observing only one of these variables, thereby bringing the method to a powerful theoretical basis. In the same years, the practical benefits of phase portraits became clear. In particular, it was used in the analysis and forecast of the animal abundance dynamics.

In this paper we propose to extend SSA to one-dimensional sequence of any type elements, including numbers, symbols, figures, etc., and, as a special case, to molecular sequence. Technically, the problem is solved almost the same algorithm as the SSA. The sequence is cut by a sliding window into fragments of a given length. Between all fragments, the matrix of Euclidean distances is calculated. This is always possible. For example, the square root from the Hamming distance between fragments is the Euclidean distance. For the resulting matrix, the principal components are calculated by the principal-coordinate method (PCo). Instead of a distance matrix one can use a matrix of any similarity/dissimilarity indexes and apply methods of multidimensional scaling (MDS). The result will always be PCs in some Euclidean space.

We called this method PCA-Seq. It is certainly an exploratory method, as its particular case SSA. For any sequence, including molecular, PCA-Seq without any additional assumptions allows to get its principal components in a numerical form and visualize them in the form of phase portraits. Long-term experience of SSA application for numerical data gives all reasons to believe that PCA-Seq will be not less useful in the analysis of non-numerical data, especially in hypothesizing.

PCA-Seq is implemented in the freely distributed Jacobi 4 package (*http://mrherrn.github.io/JACOBI4/*).

## Introduction

In the 40s of the last century, Karhunen and Loève proposed a method for processing a one-dimensional numerical time series by shifting several times and decomposition into several orthogonal time series by a multidimensional method of principal components (PCA) (Karhunen, 1947; Loève, 1948). In the 1980s Takens showed for a dynamic system, that shifts only of one observed variable allow to construct an attractor of the entire system, thereby bringing the method to a powerful theoretical basis (Takens, 1981).

The method was independently arised and applied in practice under various names (EOF, SSA, Caterpillar, etc.), including by us for the analysis of the animals abundance dynamics (Efimov, Galaktionov, 1983; Efimov et al., 1988; Efimov et al., 2003), and for other topics (Golyandina et al., 2001; Golyandina, Zhigljavsky, 2013; Golyandina et al., 2018). Today the name SSA (Singular Spectral Analysis) is most often used. The method can be extended for a sequence of any type elements, including numbers, symbols, figures, etc. and, as a special case, for molecular sequence (Efimov et al., 2018). It is the point of this article.

## Algorithm

Let there be a sequence *X* = *{x*_*1*_, *x*_*2*_, *…, x*_*N*_*}* of any type elements. Choose a lag *L, N > L > 1*. Denote by *X*_*i*_ the fragment *X* of length *L* terminated by the element *x*_*i*_, *X*_*i*_*=(x*_*i-L+1*_, *x*_*i-L+2*_, *… x*_*i-1*_, *x*_*i*_*), N ≥ i ≥ L*. Compute the matrix of Euclidean distances *D* = *(d*_*ij*_ *= d(X*_*i*_, *X*_*j*_)) between all fragments (this is always possible, for example, using the number of unmatched elements, but not only). Apply the method of principal coordinates (PCo) to the *D* and obtain its principal components PCs (Gower, 1966). Call this method <u>PCA-Seq</u>.

The usual method of finding the principal components consists in the following (Jolliffe, Cadima, 2016). Let X be a centered matrix of objects coordinates in a certain Euclidean space. We can apply to X the singular value decomposition (SVD): X = PSV^T^, where P, V^T^ – orthogonal matrices, and S – diagonal matrix of X singular values. There is possible to apply SVD to a symmetric matrix XX^T^: XX^T^ = PΛP^T^, where P – the same orthogonal matrix as for X, and Λ – a diagonal matrix of the matrix XX^T^ singular values. But XX^T^ = PSV^T^*VSP^T^ = PSSP^T^ = PS^2^P^T^. Consequently, S^2^ = Λ и S = Λ^1/2^. That is, the matrix of <u>singular</u> values of the matrix XX^T^ is the matrix of <u>eigenvalues</u> of the matrix X. Therefore, it is necessary to calculate the principal components by the formula U = PΛ^1/2^. This is very useful in practice if the number of objects is significantly less than the number of traits that are becoming more common in biological research, especially molecular ones.

More than half a century ago, Gower (Gower, 1966) found that if we calculate the matrix D of Euclidean distances between rows X, square this distances, double center and multiplication them by −1/2, then we get the XX^T^ matrix. Applying SVD to it, we get the principal components. For this reason Gower called this method the principal coordinates (PCo) analysis. However, it follows from the results of Gower that the matrix X itself is not needed and may not even exist in numerical form. To calculate the principal components of a certain set of objects, it is enough to have a matrix of Euclidean distances between them obtained by any way. If we calculate the Euclidean matrix of distances between the rows of the matrix of principal components, then it will coincide with the initial matrix of Euclidean distances D. This property can be used to verify the calculations.

PCo is quite often used for dissimilarity matrices, for which it is unknown whether they are Euclidean distances between objects or not. In the case of non-Euclidean distances, some diagonal values of the matrix Λ will be negative. Small negative diagonal values can sometimes arise due to the accumulation of computational errors. All such “components”, as well as zero ones, should be excluded from consideration.

Instead of a distance matrix one can use a matrix of any coefficients of similarity/dissimilarity. In this case it is necessary to apply methods of multidimensional scaling (MDS). The results will always be the PCs in some Euclidean space (Gower, 1966). (PCo has another name: metric multidimensional scaling with abbreviations MMDS or simply MDS. It is more correct to call MDS all methods of multidimensional scaling, but MMDS apply to PCo only.)

## Data

The amino acid sequence of the *Homo sapiens Cytb* gene was used (Q0ZFD6_HUMAN, Swiss Model repository) (table 1). The sequence (length N=380) contains two chains in positions 19−204, 259−359 and nine transmembrane helices in positions 30−56, 77−98, 113−133, 140−158, 178−200, 229−246, 288−308, 320−339, 345−372 (https://swissmodel.expasy.org/repository/uniprot/Q0ZFD6).

**Table 1.** The amino acid sequence of the *Homo sapiens Cytb* gene (Q0ZFD6_HUMAN, Swiss Model repository, https://swissmodel.expasy.org/repository/uniprot/Q0ZFD6). The top line is the sequence identifier, lower − the sequence itself. The first and last fragments of length 8 are highlighted in bold type (see Table 2.).

|  |
| --- |
| >tr\|Q0ZFD6\|Q0ZFD6_HUMAN Cytochrome b OS=Homo sapiens GN=CYB |
| **MTPMRKTN**PLMKLINHSFIDLPTPSNISAWWNFGSLLGACLILQITTGLFLAMH<br>YSPDASTAFSSIAHITRDVNYGWIIRYLHANGASMFFICLFLHIGRGLYYGSFLYS<br>ETWNIGIILLLATMATAFMGYVLPWGQMSFWGATVITNLLSAIPYIGTDLVQWI<br>WGGYSVDSPTLTRFFTFHFILPFIIAALATLHLLFLHETGSNNPLGITSHSDKITFH<br>PYYTIKDALGLLLFLLSLMTLTLFSPDLLGDPDNYTLANPLNTPPHIKPEWYFLF<br>AYTILRSVPNKLGGVLALLLSILILAMIPILHMSKQQSMMFRPLSQSLYWLLAAD<br>LLILTWIGGQPVSYPFTIIGQVASVLYFTTILILMPT**ISLIENKMLKWA** |

## Processing

Denote by N_L_ = N-L+1. For L = 2, …, 24, the sequence Q0ZFD6 represented as Seq_L_ matrix of size N_L_*L (Table 2 with L = 8, as an example). For each Seq_L_, the matrix A_L_ of size N_L_*20 – the each amino acid content in the fragment and the H_L_ vector of length N_L_ – the fraction of fragment positions coinciding with transmembrane helices are additionally calculated. For all matrices Seq_L_, matrices of Euclidean distances between fragments are calculated (square root of the *p*-distance (Hamming distance) between couple of fragments is used as the Euclidean distance (Efimov et al., 2013)).

**Table 2.**
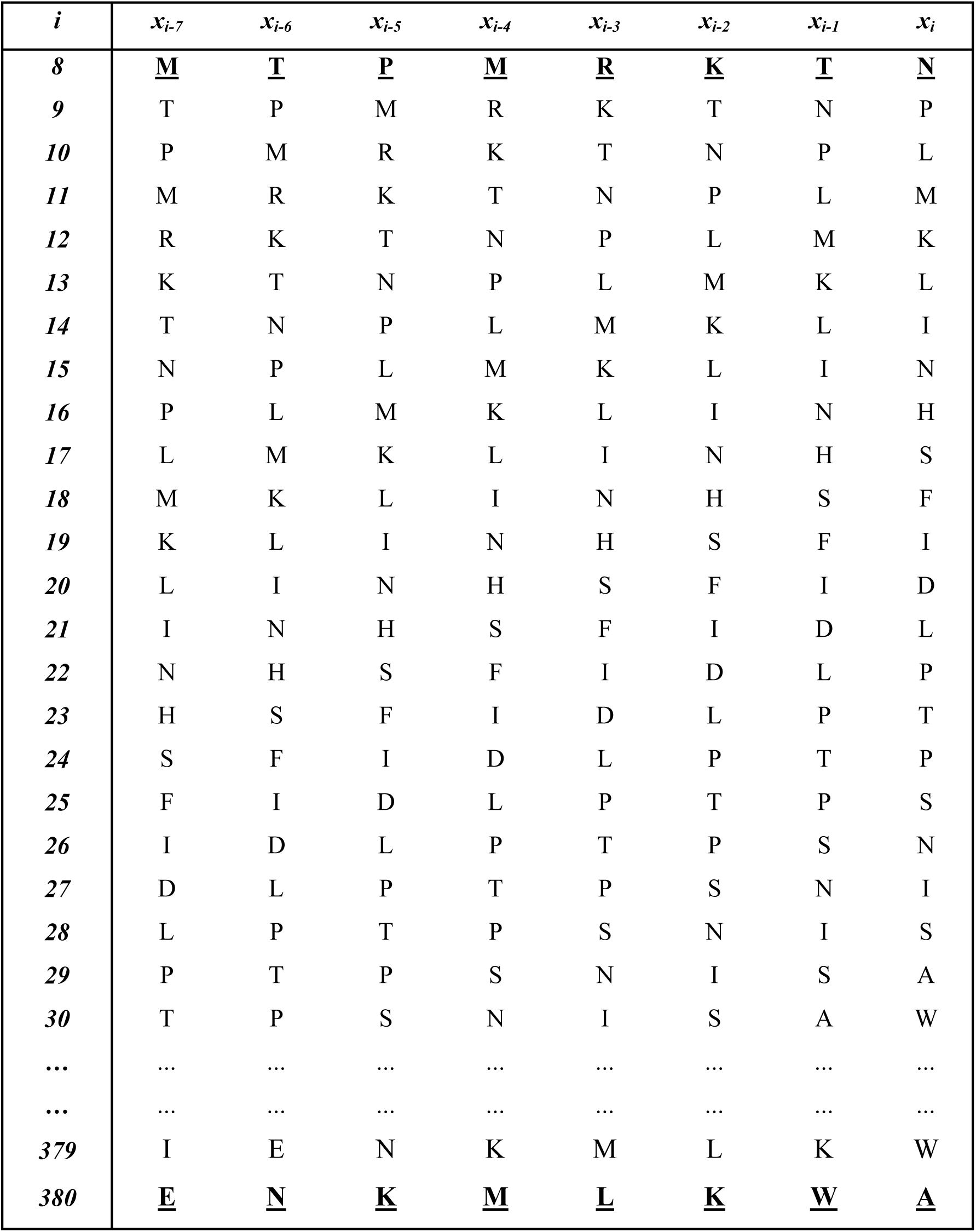
Takens embedding transformation of the amino acid sequence from Table 1. Its first and last rows are first and last fragments of the sequence, the same as in Table 1. Number of a fragment is defined by number of its last amino acid therefore the table begins with row 8.

For all matrices of Euclidean distances, its PCs are calculated by the method of PCo (Gower, 1966). The matrices PC-Seq_L_, A_L_ and the vector H_L_ were combined into one matrix, and for it, the matrix of Pearson correlation coefficients was calculated between all columns. The correlation coefficients exceeding on the threshold 0.316 by module (r^2^ ∼ 0.1, i.d. 10%; p < 10^-8^) were considered only. Jacobi 4 package was used for calculations (Polunin et al., 2014).

## Results

For the first principal component PC1-Seq_L_, the correlation exceeding the threshold was found with a fraction of helix positions (0.370 ⩽ r ⩽ 0.547 in the range 4 ⩽ L ⩽ 18; r_max_ = 0.547 for L = 12), leucine content in the fragment (0.95 ≤ in the range 2 ≤ L ≤ 17; r_12_ = 0.974), proline content (0.331 ≤ r ≤ 0.364 in the range 9 ≤ L ≤ 14) and tyrosine content (0.318 ≤ r ≤ 0.351 in the range 14 ≤ L ≤ 18), what is more, correlations PC1<u>-</u> Seq_L_ with the contents of proline and tyrosine have inverse sign in relation of correlations with the fraction of helix positions and the leucine content in the fragments. Graph of PC1<u>-</u>Seq_L_ against the background of the fraction of helix positions is shown in Fig. 1, the dynamics of the leucine content against the same background on fig. 2, the scatterplot of PC1-Seq_L_ vs the leucine content in the fragment is shown in fig. 3 (all for L = 12).

**Fig. 1.**
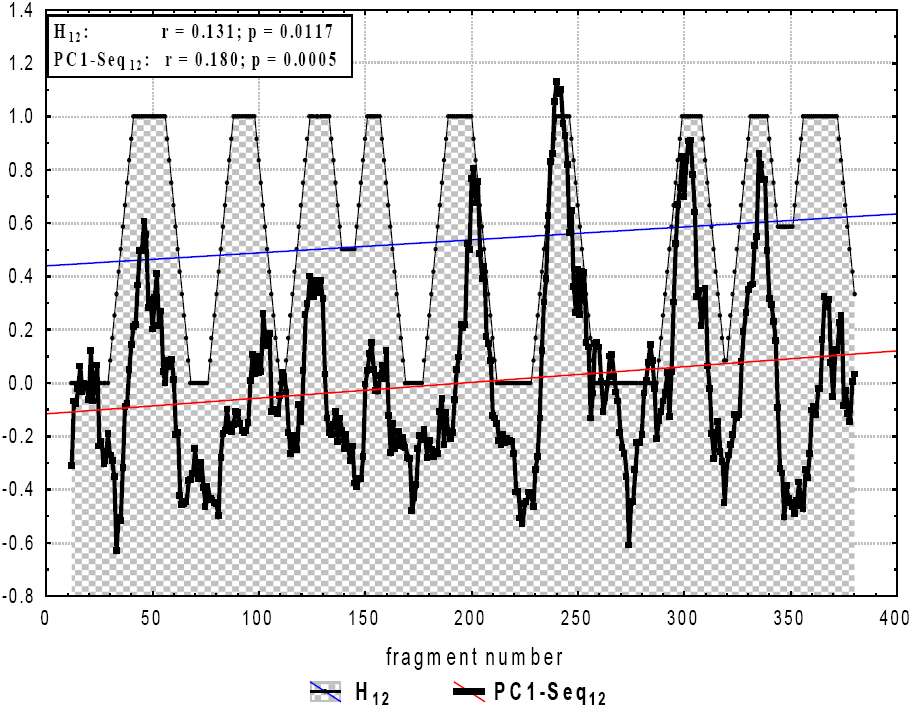
PC1-Seq_12_ − first principal component of *Homo sapiens Cytb* gene sequence (Q0ZFD6_HUMAN, Swiss Model repository) and H_12_ − fraction of helix position in sequence fragments of length L = 12 (r = 0.547; N = 369; p < 10^-9^).

**Fig. 2.**
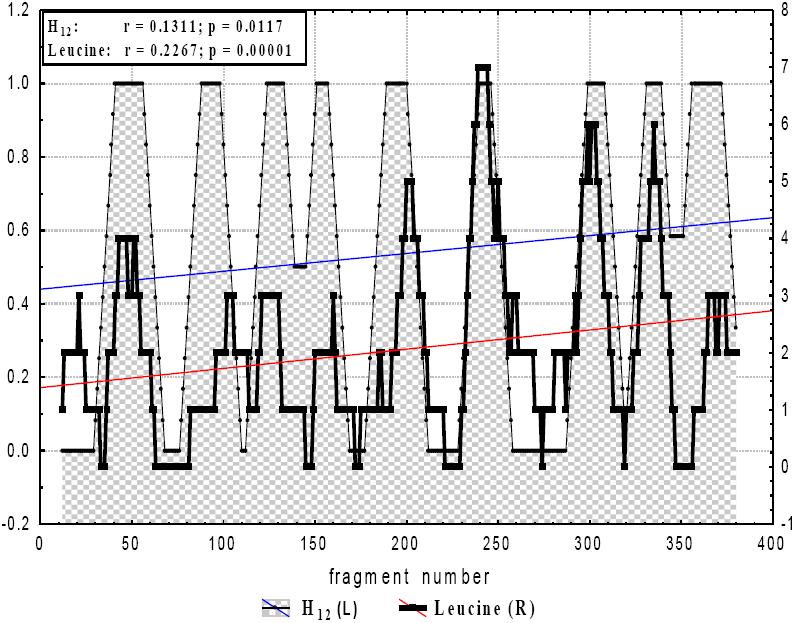
Content of Leucine in *Homo sapiens Cytb* gene sequence fragments (Q0ZFD6_HUMAN, Swiss Model repository) and H_12_ − fraction of helix position in sequence fragments of length L = 12 (r = 0.499; N = 369; p < 10^-9^).

**Fig. 3.**
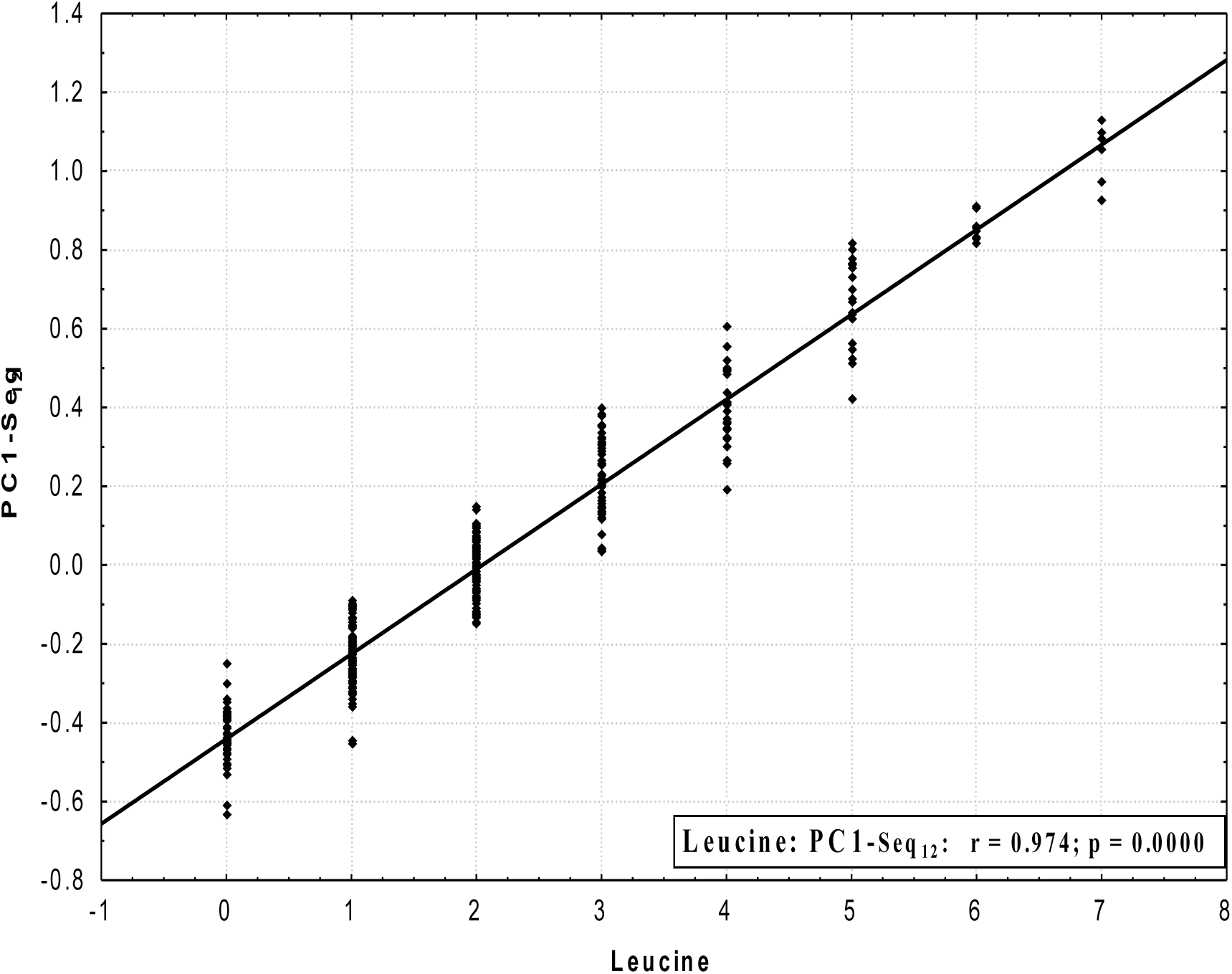
Content of Leucine in *Homo sapiens Cytb* gene sequence fragments (Q0ZFD6_HUMAN, Swiss Model repository) vs first principal component PC1-Seq_12_ (L=12; r = 0.974; N = 369; p < 10^-9^).

## Discussion

Note that we did not specifically look for any information about the amino acids content in the fragments and about second structure of sequence. The only thing that we investigated is how much the fragments coincide with each other by amino acids in total for all L positions. If we had set another measure of similarity, then perhaps we would have discovered some other regularity. In this case, this one is found.

PCA-Seq is certainly an exploratory method, as its particular case SSA. For any sequence, including molecular, PCA-Seq without any additional assumptions allows to get its principal components in a numerical form and visualize them in the form of phase portraits. Today SSA for numerical series is a huge scientific field with applications in the various sciences. There is no doubt that the analysis of non-numeric sequences will be scientific field, no less in scope than SSA.

It should be noted that the approach of calculation through a covariation (correlation) matrix is used in the standard SSA only, and the MDS methods, including PCo, despite a more than half a century history, are almost unknown. This gives reason to hope that PCA-Seq can be useful in the analysis of real data, especially in hypothesizing.

PCA-Seq is a particular case of geometric approach, in which any similarity/ dissimilarity relations between objects is modeled by the distance between points in a certain Euclidean space. In this case, the objects are fragments of any sequence by length L, including non-numeric, in particular molecular one. Orthogonal rotations of the entire set of points remain unchanged the distances between them. This allows to calculate such axes, in the projection on which the maximum variances of the set of points are reached. These axes always exist, exactly they are the principal components. By construction, they are not statistically correlated with each other. This does not mean at all that they are meaningfully independent.

In particular, for a time series it is a general rule that their PCs, despite being uncorrelated, break up into couples in which one component is derivative of another. When one component shifts from another by a quarter of a period, the correlation appears again. In successful cases, this allows to predict the future values of one component from the already known values of another and, thereby, to some extent predict the initial series. The sine – cosine couple is a bright example. Moreover, it is possible that there is a third component, as a rule, a trend, which, despite the lack of correlation with the first two, modulates them by amplitude. Thereby, it is a part of the general interconnected complex. In 3D phase space such components form a funnel.

PCA-Seq is implemented in the freely distributed Jacobi4 package (http://mrherrn.github.io/JACOBI4/).

## Conclusion

PCA-Seq is promising for processing molecular sequences, but not only.

## Acknowledgements

Supported by Russian Foundation for Basic Research (#19-07-00658).

The authors are grateful to D.A. Afonnikov, P.N. Menshanov and two anonymous reviewers for useful discussion and constructive comments.

